## supplemental table.1 for "IL-6 is a Key Factor in the Formation of Gut Tissue Resident Memory T Cells from Naïve T cells"

**Supplemental Table 1. List of antibodies used in the study.**

| Target | Fluorophore | Clone | Company | Cat. Number | |
|---|---|---|---|---|---|
| CD103 | PE-cy7 | Ber-ACT8 | Biolegend | 350212 | FACS |
| CD69 | BV480 | FN50 | BD | 747519 | FACS |
| CCR9 | APC | L053E8 | Biolegend | 358908 | FACS |
| CCR5 | BV421 | J418F1 | Biolegend | 359118 | FACS |
| CD3 | BV605 | UCHT1 | Biolegend | 3004660 | FACS |
| CD4 | AF700 | RPA-T4 | Biolegend | IV T114 | FACS |
| CD8 | BB700 | RPA-T8 | BD | 566452 | FACS |
| CD45RO | BV650 | UCHL1 | BD | 748367 | FACS |
| β7 | PE | FIB504 | BD | 555945 | FACS |
| β7 | APC | FIB504 | BD | 551082 | FACS |
| β1 (CD29) | PE | MAR4 | BD | 555443 | FACS |
| α4β7 | FITC | ACT-1 | In lab | - | FACS |
| STAT1 | - | 1/Stat1 | BD | 610116 | WB |
| p-STAT1 | - | 4a | BD | 612233 | WB |
| p-STAT2 | - | | Millipore | 07224 | WB |
| STAT3 | - | D1B2J | Cell Signal | 30835 | WB |
| p-STAT3 | - | D3A7 | Cell Signal | 9145 | WB |
| Actin | - | AC-15 | Santa Cruz | sc-69879 | WB |
